# The State of Software in Evolutionary Biology

**DOI:** 10.1101/031930

**Authors:** Diego Darriba, Tomáš Flouri, Alexandros Stamatakis

## Abstract

With Next Generation Sequencing Data (NGS) coming off age and being routinely used, evolutionary biology is transforming into a data-driven science.

As a consequence, researchers have to rely on a growing number of increasingly complex software. All widely used tools in our field have grown considerably, in terms of the number of features as well as lines of code. In addition, analysis pipelines now include substantially more components than 5-10 years ago.

A topic that has received little attention in this context is the code quality of widely used codes. Unfortunately, the majority of users tend to blindly trust software and the results it produces. To this end, we assessed the code quality of 15 highly cited tools (e.g., MrBayes, MAFFT, SweepFinder etc.) from the broader area of evolutionary biology that are used in current data analysis pipelines.

We also discuss widely unknown problems associated with floating point arithmetics for representing real numbers on computer systems. Since, the software quality of the tools we analyzed is rather mediocre, we provide a list of best practices for improving the quality of existing tools, but also list techniques that can be deployed for developing reliable, high quality scientific software from scratch.

Finally, we also discuss journal and science policy as well as funding issues that need to be addressed for improving software quality as well as ensuring support for developing new and maintaining existing software.

Our intention is to raise the awareness of the community regarding software quality issues and to emphasize the substantial lack of funding for scientific software development.

---

With Next Generation Sequencing Data (NGS) coming off age and being routinely used, it cannot be disputed that evolutionary biology is becoming even more quantitative. With massive amounts of data there is also a paradigm shift from a hypothesis-driven to a data-driven science, irrespective of one’s own philosophical perceptions of whether this represents a positive or negative development.

Our field is also becoming a true computational science which routinely relies on supercomputers (e.g., Misof et al. (2014) or Jarvis et al. (2014)). This is a transition other disciplines such as astrophysics or fluid dynamics accomplished decades ago.

The common denominator of the above trends is that researchers have to rely on a larger number of increasingly complex software. By software complexity we refer to the fact that all widely used tools have grown considerably, in terms of the number of features as well as lines of code. For instance, MrBayes (Ronquist et al. 2012) had approximately 49,000 lines of code in 2005 and about 94,000 in 2014. Phylogenetic inference software now supports a substantially larger set of models (e.g., substitution models), hardware platforms (e.g., GPUs, clusters, etc.), and types of parallelism (e.g., fine-grain, coarse-grain, hybrid approaches).

In addition, software complexity can also be quantified by means of the component count in current analysis pipelines. In the ‘Sanger days’, the analysis pipeline was rather straightforward, once the sequences were available. For a phylogenetic study it consisted of the following steps: align → infer tree → visualize tree. For NGS data and huge phylogenomic datasets, such as the insect transcriptome (Misof et al. 2014) or bird genome evolution (Jarvis et al. 2014) projects, pipelines have become substantially longer and more complex. They also require user expertise in an increasing number of bioinformatics areas (e.g., orthology assignment, read assembly, dataset assembly, partitioning of datasets, divergence times inference, etc.). In addition, these pipelines require a plethora of helper scripts to transform formats, partially automate the workflow, and connect the components. Helper scripts are typically written in languages such as `perl` (a language that is highly susceptible to coding errors due to lack of typing) or `python` that uses dynamic typing and can thus not be subjected to a comprehensive type-check either. The term ‘typing’ refers to the data types of variables (e.g., integer or floating point) that are passed to and returned by functions. Without strict typing a function expecting an integer argument can be invoked with a floating point value as an argument and exhibit undefined or unexpected behavior.

Our main concern is that, if each code (henceforth, we use code as synonym for software) or script component i used in such a pipeline has a probability of being ‘buggy’ *P_i_*, the probability that there is a bug in the pipeline increases dramatically with the number of components. If detected too late, errors in the early stages of pipelines (e.g., NGS assembly) for large-scale data analysis projects can have a dramatic impact on downstream analyses such as phylogenetic inferences or dating. They will all have to be repeated. In fact, this has happened in every large-scale data analysis project we have been involved in thus far. Given that our field needs to compete with established computational sciences for scarce supercomputing or cloud resources, repeating large phylogenomic analyses can result in a substantial waste of computational resources.

Another concern is that evolutionary analysis software is frequently used as a black box with default parameters and without a proper understanding of the underlying theory or algorithms. Given the large set of tools modern evolutionary biologists need to deploy to ‘get a paper published’, this user behavior is nonetheless understandable. There is an evident trade-off between the thoroughness of computational analyses and the publication rate. While this reality is difficult to change, the issue should be addressed at the teaching level. Our perception is that graduate and undergraduate training in biology needs to become substantially more quantitative.

Based on the prolegomena, our goals in this paper are to assess the quality of current software and to propose potential solutions, including software analysis tools, for improving the quality of evolutionary biology software. We wish to emphasize that the quality measures we deploy only represent one option for assessing software. Software quality is not necessarily an indicator for correctness, but a correlation does exist (e.g., Briand et al. (1999, 2000)).

For assessing software quality we downloaded and scrutinized-using a common set of criteria-15 frequently used and cited codes that often form the basis of data analyses published in *Systematic Biology* and related journals. For comparison, we also analyzed an Astrophysics code developed at our research institute because Astrophysics is a more mature computational science discipline. Based on the software analysis results, we provide our personal and subjective list of best practices and discuss some science policy issues that need to be addressed for improving software quality and for supporting scientific software development.

Note that, it is absolutely not our intention to criticize any of the authors and developers of the codes we assessed. They have all made major contributions to the field. We also need to keep in mind that a large fraction of the developers has never received formal training in computer science and that they are mostly self-taught programmers. Moreover, it is quite typical that the careers of PIs in bioinformatics are based on one or more widely used tools they have developed. As they become more senior and manage larger research groups, there is less time available to maintain and occasionally re-design the tools, despite the fact that they know how to implement software ‘the right way’ in principle. In addition, they are mostly reluctant to delegate this task to graduate students or postdocs because they should work on more interesting projects instead of merely re-engineering widely used software.

Given that most software for evolutionary biology is distributed under the GNU GPL license, users and critics should keep the following quote from the GNU GPL license in mind: “The copyright holders and/or other parties provide the program ‘as is’ without warranty of any kind, either expressed or implied, including, but not limited to, the implied warranties of merchantability and fitness for a particular purpose. The entire risk as to the quality and performance of the program is with you. Should the program prove defective, you assume the cost of all necessary servicing, repair or correction.”

Thus, our goal is to emphasize that users should be aware of the fact that software is imperfect. Furthermore, because of the increasing reliance on software in current day biology, there exists a substantial funding, sustainability, and maintenance issue that needs to be addressed.

## Software & Analysis Methods

### Software

We selected highly cited open-source tools from the following areas: phylogenetic inference, population genetics, multiple sequence alignment, divergence time estimation, multi-species coalescence, sequence simulation, and de novo assembly. Note that tools from all of these areas can be used in evolutionary biology data analysis pipelines. We deliberately omitted codes from our lab in this list, to avoid any potential bias; our codes are not better than the software analyzed here.

In Table 1 we list the codes we assessed in each domain.

**Table 1:**
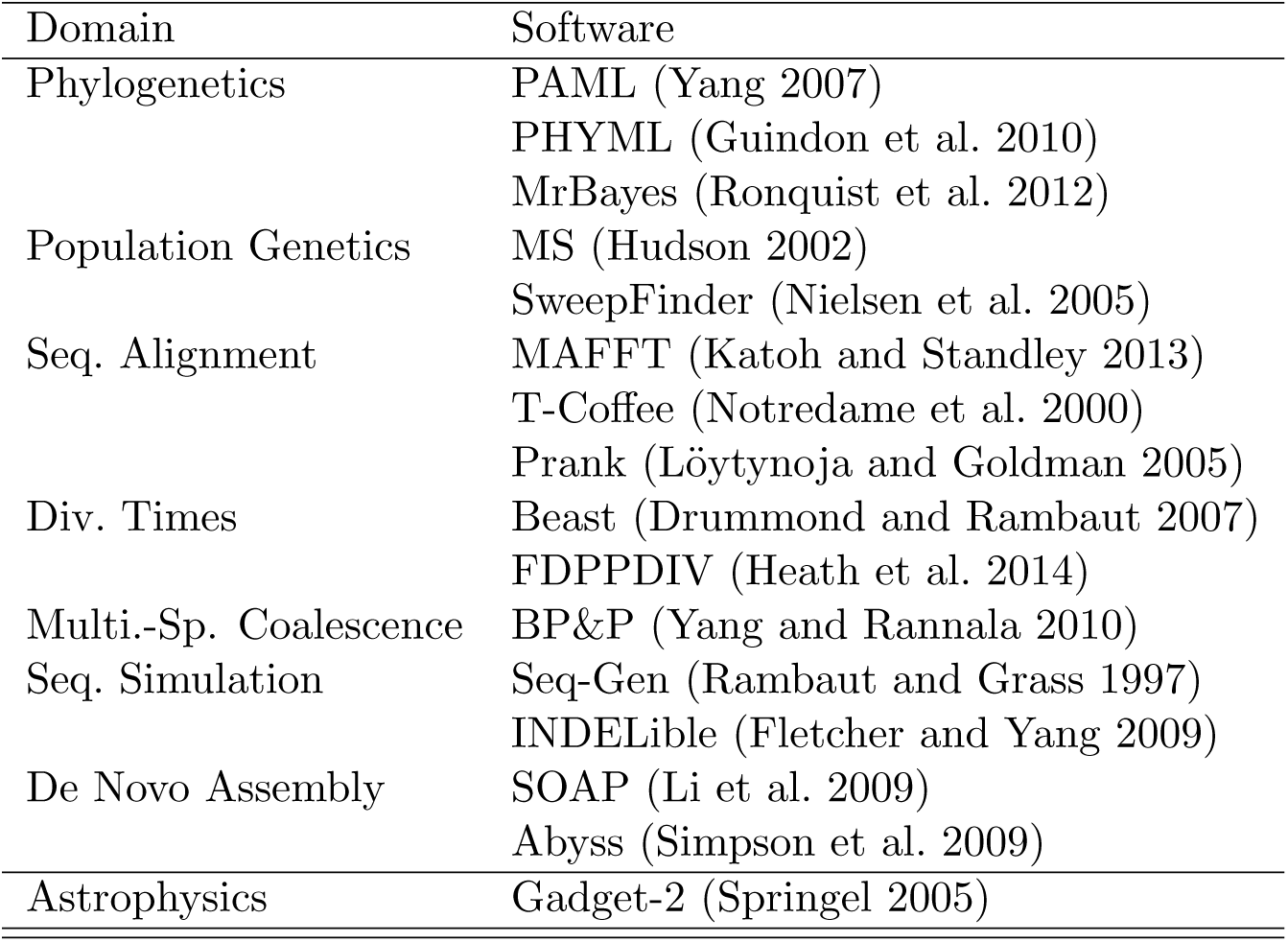
Evaluated software packages per application domain

### Code Analysis Criteria

Since we analyzed a comparatively large number of codes, we deployed rather simple and straightforward techniques to analyze them.

Initially, we compiled all codes using the standard GNU compilers (gcc/g++) as well as the clang compiler by Apple. We enabled all reasonable warning flags in the two C/C++ compilers as well as analogous flags in JAVA for analyzing BEAST (see supplement for details). We classified GNU compiler warnings into major warnings that are potentially dangerous and minor warnings that are less dangerous, but should be fixed nonetheless (see supplement for the classification criteria). We count and classify compiler warnings, because we assume that the more warnings a code produces, the more likely it is to behave in an unexpected way. However, this does not automatically mean that the results computed by these codes are incorrect, since a code that produces no warnings can yield incorrect results.

Then, we executed the codes using the `valgrind` tool (http://valgrind.org/) to detect potential memory leaks, illegal memory accesses, lost memory blocks, etc. We classified results into three categories: ‘clean’ when running the codes with `valgrind` did not generate any warnings, ‘invalid’ for read or write accesses at an invalid RAM address, or ‘leaks’ when allocated memory was not properly freed again. Memory errors or incorrect usage of memory serves as an indicator for the probability of crashes or unspecified behavior, when accessing values at invalid or uninitialized RAM locations.

Thereafter, we used the `grep` text searching tool to identify a typical programming error associated with the `C malloc()` routine that is used to allocate a memory block of *n* bytes in RAM. Frequently, this function is invoked with integer data types that are too small for representing *n* to allocate large chunks of memory. In our analyses, we distinguish between three `malloc()` usage errors: ‘NoCast’ (i.e., missing typecast) and ‘MisCast’ (misplaced cast) and ‘WrongCast’ (incorrect cast). For the `new[]` operator in C++ we use an analogous classification. Examples for these error types (e.g., in MrBayes and ms) are provided in the supplement. While for smaller datasets this incorrect usage will have no effect, programs are likely to crash when deployed for analyzing NGS datasets on powerful multi-core servers which are nowadays often equipped with 128 or 256GB RAM.

Another code feature that we consider as being important is the use of so-called assertions (e.g., the `assert()` function in C, see supplement for a classic `assert()` example). We assessed the usage of assertions by calculating the number of assertions per 1000 lines of code. Assertions contain logical clauses about variables that must be true when the program conducts an assertion call, otherwise the program fails. The use of assertions is associated with code correctness. In theoretical computer science, there exists a framework, the so-called Hoare logic (Hoare 1969), for proving program correctness. It works by inserting assertions (Boolean statements about variable states) at appropriate positions in the code and proving that they will never fail. While proving the correctness of the complex scientific codes we scrutinize here using Hoare logic is not feasible, we consider that the frequent use of assertions in a program is an indicator of code quality. Note that, PHYML uses assertion-like conditional `if`-statements instead. We discuss why we do not think that this is good practice in the supplement.

To obtain a rough estimate of code complexity, we also counted the lines of code (LoC) in each of the programs using the `cloc` (http://cloc.sourceforge.net/) command that excludes comments and empty lines. For some programs we also generated histograms that illustrate code growth over the last years (see supplement). The LoC metric of course does not directly reflect code complexity, but can serve as a rough proxy for it.

A more elaborate criterion for assessing code complexity is the degree of code duplication, that is, how many copies of identical code are present in the source files. In general, code duplication represents a bad programming practice. If a bug is detected and fixed in one copy of the duplicated code, it needs to be fixed in all duplicates. Mostly, these duplicates are not properly documented and potentially difficult to find. Thus, software with a high degree of code duplication is more difficult to maintain and thus more likely to contain errors.

Overall, the above criteria have been selected (i) because they are easy to apply to a large number of diverse codes and because (ii) there exists a correlation (e.g., Briand et al. (1999, 2000)) between quality and the probability of erroneous program behavior, that is, crashes or calculation of incorrect results.

## Software Analysis Results

A detailed analysis of the codes, including appropriate source code examples, is provided in the on-line supplement.

We summarize the results of our standard tests in Table 2 for all PAML components individually and in Table 3 for all other programs including the PAML core code. The results obtained by the Simian tool (http://www.harukizaemon.com/simian/) that quantify the degree of code duplication are summarized in Table 4.

One general observation is that the clang compiler issues substantially more warnings than the GNU compilers. This is because it performs a so-called static code analysis, that is, a more thorough check, including stricter type checking. Another general trend is the infrequent use of assertions as well as rather sloppy memory management. While memory leaks can be harmless, invalid memory accesses (Prank, MrBayes, MAFFT) are more likely to yield unspecified behavior.

**Table 2:**
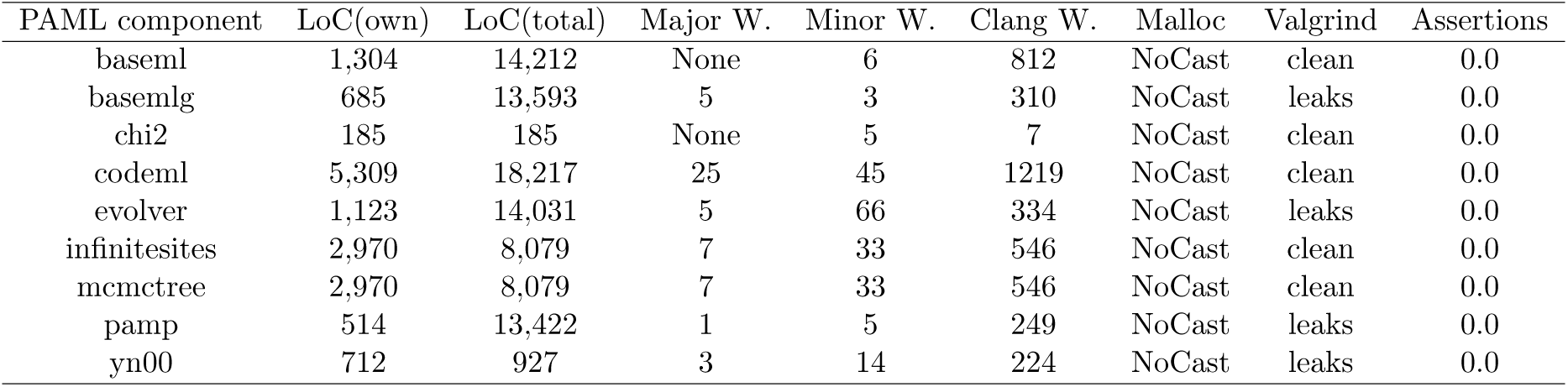
PAML components. LoC(own) is the number of effective lines of code that belong only to the component. LoC(total) is the total number of effective lines of code for each component, including code shared with other components. Columns ‘Major W.’ and ‘Minor W.’ give the major and minor GNU compiler warnings and ‘Clang W.’ reports the number of clang warnings. Column ‘Malloc’ provides the malloc() casting error, ‘Valgrind’ the memory behavior and ‘Assertions’ the number of assertions per 1000 lines of code.

**Table 3:**
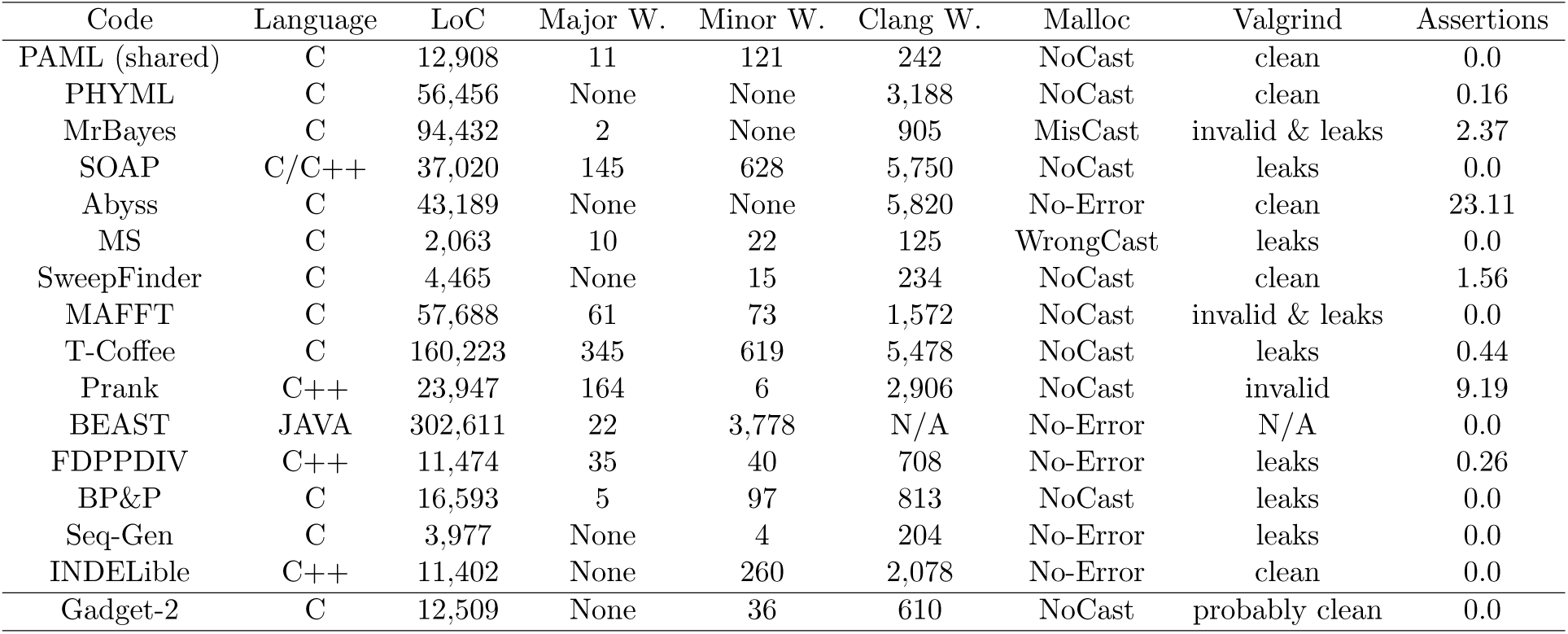
PAML values refer to parts of the source code that is shared among all individual components of Table 2 Column ‘Language’ denotes the programming language and column ‘LoC’ is the total number of effective lines of code. Columns ‘Major W.’ and ‘Minor W.’ give the major and minor GNU compiler warnings and ‘Clang W.’ reports the number of clang warnings. Column ‘Malloc’ provides the malloc() casting error, ‘Valgrind’ the memory behavior. We denote the Gadget-2 code as ‘probably clean’ since we interrupted the valgrind analysis that did not report any errors after 30 minutes of run-time. Finally, column ‘Assertions’ represents the number of assertions per 1000 lines of code.

| Code | Language | LoC | Major W. | Minor W. | Clang W. | Malloc | Valgrind | Assertions |
| --- | --- | --- | --- | --- | --- | --- | --- | --- |
| PAML (shared) | C | 12,908 | 11 | 121 | 242 | NoCast | clean | 0.0 |
| PHYML | C | 56,456 | None | None | 3,188 | NoCast | clean | 0.16 |
| MrBayes | C | 94,432 | 2 | None | 905 | MisCast | invalid & leaks | 2.37 |
| SOAP | C/C++ | 37,020 | 145 | 628 | 5,750 | NoCast | leaks | 0.0 |
| Abyss | C | 43,189 | None | None | 5,820 | No-Error | clean | 23.11 |
| MS | C | 2,063 | 10 | 22 | 125 | WrongCast | leaks | 0.0 |
| SweepFinder | C | 4,465 | None | 15 | 234 | NoCast | clean | 1.56 |
| MAFFT | C | 57,688 | 61 | 73 | 1,572 | NoCast | invalid & leaks | 0.0 |
| T-Coffee | C | 160,223 | 345 | 619 | 5,478 | NoCast | leaks | 0.44 |
| Prank | C++ | 23,947 | 164 | 6 | 2,906 | NoCast | invalid | 9.19 |
| BEAST | JAVA | 302,611 | 22 | 3,778 | N/A | No-Error | N/A | 0.0 |
| FDPPDIV | C++ | 11,474 | 35 | 40 | 708 | No-Error | leaks | 0.26 |
| BP&P | C | 16,593 | 5 | 97 | 813 | NoCast | leaks | 0.0 |
| Seq-Gen | C | 3,977 | None | 4 | 204 | No-Error | leaks | 0.0 |
| INDELible | C++ | 11,402 | None | 260 | 2,078 | No-Error | clean | 0.0 |
| Gadget-2 | C | 12,509 | None | 36 | 610 | NoCast | probably clean | 0.0 |

**Table 4:** Results of a code duplication analysis using the Simian tool. The column ‘Lines checked’ refers to the total number of source lines and ‘Files checked’ to the total number of source files analyzed with Simian. Note that, the ‘Lines checked’ number is not identical to the LoC numbers reported in T++++++ables 2 and 3, since the Simian tool does not take header files into account. Column ‘Duplicate lines’ provides the number of duplicate lines detected and ‘Blocks’ provides the total number of contiguous duplicated blocks of code. Finally, column ‘Files’ gives the number of files in which duplicated code was detected.

| Code | Lines checked | Files checked | Duplicate lines | Blocks | Files |
| --- | --- | --- | --- | --- | --- |
| PAML | 22,200 | 17 | 1,210 | 120 | 11 |
| PHYML | 42,786 | 73 | 5,878 | 549 | 32 |
| MrBayes | 70,680 | 19 | 21,862 | 1,680 | 10 |
| SOAP | 27,514 | 116 | 10,107 | 527 | 72 |
| Abyss | 37,038 | 212 | 4,245 | 441 | 71 |
| MS | 1,718 | 24 | 186 | 21 | 9 |
| SweepFinder | 3,777 | 12 | 293 | 28 | 3 |
| MAFFT | 45,045 | 72 | 28,630 | 1,647 | 59 |
| T-Coffee | 82,758 | 196 | 19,345 | 1,325 | 58 |
| Prank | 16,124 | 67 | 5,318 | 462 | 43 |
| BEAST | 228,316 | 2,336 | 64,024 | 4,786 | 1,151 |
| BP&P | 14,332 | 5 | 502 | 56 | 3 |
| Seq-Gen | 3,244 | 44 | 206 | 25 | 6 |
| INDELible | 9,840 | 7 | 1,954 | 106 | 5 |
| Gadget-2 | 9,770 | 31 | 3,314 | 180 | 31 |

We also observe a high degree of code duplication in some codes (e.g., MrBayes, SOAP, MAFFT, Prank, BEAST).

Overall, the perfect software does not seem to exist, with the exception of Abyss maybe, if we ignore the `clang` warnings. The Astrophysics code is not perfect either (e.g., using no assertions at all), despite the fact that it comes from a more traditional field of computational science. However, our set of criteria allows to rapidly identify potential problems that can, in most cases easily be fixed.

## Discussion

We have scrutinized 15 widely used codes for evolutionary data analyses using a simple set of tools and criteria that can be deployed to improve code quality, even without understanding the source code. Evidently, software quality can only be assessed with open-source codes, hence we strongly argue in favor of open-source such that users at least have a chance to assess code quality.

We have detected several errors that are common to almost all tools and that are comparatively easy to fix. Again, we do not intend to criticize the authors of the tools, given their time and resource constraints with respect to extending and maintaining software. We want to emphasize that more awareness about code quality and, perhaps more importantly, worrying about correctness is necessary since the research produced by our community increasingly relies on the results produced by an entire swarm of tools in analysis pipelines.

We initially discuss some good practices for code development in the hope that they will be broadly adopted by the community and help to reduce the number of bugs. Then, we discuss issues pertaining to floating-point arithmetics and reproducibility of numerical results. Finally, we discuss funding policy issues, that is, what sort of mechanisms might be required to ensure sustainable maintenance, support, and quality improvements in scientific software.

### Best Practices

Some of our recommendations can be directly derived from the simple criteria we have deployed. Therefore, a good code should:

- be compiled with all compiler warning flags enabled using several compilers (e.g., `icc`, `clang`, `gcc`)
- should be analyzed with `valgrind` for memory leaks and invalid read/write accesses
- should be checked for `malloc()` type casting errors should use as many assertions as possible and reasonable

Although we doubt if this is feasible, it might represent a good idea to ask reviewers of software papers (e.g., the *Syst. Bio*. software track or application notes in *Bioinformatics)* to check software they review according to the above straightforward criteria. Alternatively, journals could impose upon authors that the codes they submit for publication need to be compiled and checked accordingly prior to submission. This could be implemented by asking authors to provide appropriate code quality transcripts. Finally, One should also put special emphasis on software quality issues (e.g., no `clang` warnings, usage of assertions, checks with `valgrind`) when teaching programming practicals at the graduate and undergraduate level.

Assertions are also particularly useful for debugging, since users often provide incomplete bug reports. In contrast to this, when an assertion fails, users will typically report the failed assertion including the source file name and the line in the code which substantially accelerates problem identification. Also, assertions are the only mechanism we considered that helps to partially assess code correctness and not only identify potential programming errors.

While invalid read/write access need to be fixed, memory leaks, in particular when programs do not free all the memory they use (e.g., several PAML components) once they terminate should be fixed. Such program termination leaks may become problematic when one intends to integrate leaky code as a library component into some larger project. Unfortunately, it is always hard to predict which software one writes will become widely used and how much effort should be spent on code quality.

The above best practices can be easily applied without investing too much effort and will certainly improve code quality as well as help to reduce the number of implementation-induced bugs. Evidently, we also need to worry about conceptual errors that affect correctness, such as the for a long time undetected error in Hastings ratio calculations (Holder et al. 2005) in Bayesian inference programs.

Another question is what else *could* be done to improve code quality in an ideal setting. Users often tend to forget that many codes, specifically in population genetics and phylogenetics, use statistical models defined on real numbers. As a consequence, they are at the mercy of floating point arithmetics with round-off errors and numerical under- or overflows. Therefore, every programmer in this area should read the classic paper “What Every Computer Scientist Should Know About Floating Point Arithmetic” by Goldberg (1991). The most important thing to know is that in floating point arithmetics associativity (i.e., (*x* + (*y* + *z*)) = ((*x* + *y*) + *z*)) does not necessarily hold because of round-off errors. Note that, the order of arithmetic operations and thus the degree of deviations due to round-off errors depends on (i) the compiler used (ii) the hardware features that are being used, and (iii) on how the programmer orders the arithmetic operations. Therefore, different ML program implementations (e.g., RAxML and PHYML) can yield different log likelihood scores.

However, even the same program can return different values when the likelihood calculations are parallelized over sites, depending on the number of processors being used. Thus, different numbers of processors can yield different tree topologies and, as a consequence, ML inference results may not be reproducible. For instance, we executed the AVX version of RAxML twice (data available at https://github.com/stamatak/softwareQuality), once in the sequential version and once with the PThreads version as follows:

~~~
raxmlHPC-AVX -p 12345 -m GTRGAMMA -s 354 -n T1
raxmlHPC-PTHREADS-AVX -T 2 -p 12345 -m GTRGAMMA -s 354 -n T2
~~~

The only difference between the two calls is that the addition order of per-site log likelihoods and per-site derivatives for optimizing branch lengths is changed due to the parallelization. The dataset we used is a single-gene alignment of 354 ITS sequences with 460 sites (Grimm et al. 2006) that was known to have a ‘rough’ likelihood surface. In other words, it exhibits numerous local maxima that cannot be distinguished from each other using statistical significance tests. Simply because the numerical deviations make the tree searches follow distinct paths, the two, in theory identical invocations, yield different final trees with log likelihood scores of -6562.158295 versus -6562.158171 and a relative Robinson-Foulds distance (Robinson and Foulds 1981) of 8.26%. Of course, any likelihood-based significance test comparing the two trees shows that they are not significantly different from each other.

As a consequence, in an ideal world we should also carry out a theoretical round-off error analysis for our codes. As shown above, this is particularly critical for ML codes that strive to obtain a single point estimate. Numerical issues are far less problematic for Bayesian inferences because they sample a distribution. In the supplement we also provide an example of how so-called de-normalized floating point values can affect program performance.

Finally, since the issue of software quality is just emerging, it might be extremely helpful to consult with software engineering experts. In addition, there already exists a plethora of tools that can assess the quality of the given software architecture and more advanced tools for explicitly finding bugs.

For instance, there is the `pmccabe` tool for assessing function complexity in C and C++ codes (https://people.debian.org/~bame/pmccabe/). For this, it calculates the so-called McCabe cyclomatic complexity (McCabe 1976) of functions. Typically, when the complexity of a function exceeds a score of 10 or 15 the function should be split into several sub-modules. A quick analysis of the main RAxML source file `axml.c` with the following command `pmccabe -f axml.c` revealed that in this source file alone there are 22 functions with a cyclomatic complexity score that exceeds 15.

Furthermore, static code analysis tools analogous to the seminal Lint (Johnson 1977) tool should be deployed. The `clang` compiler partially does this. As described in the supplement, FindBugs (http://findbugs.sourceforge.net) can be used for Java codes such as BEAST. Code duplication identification tools such as Simian should also be routinely used during code development. Finally, we recommend use of code coverage tools that identify code that will never be executed.

Another major method for improving code quality and being more confident about correctness is testing, such as unit tests or integration tests. There is a vast amount of research on, and methods for, software testing. A good starting point is the book on the art of software testing by Myers et al. (2011). The current testing practice in our field appears to be that testing is mostly delegated to users.

Thus, for programmers, we further recommend the following best practices:

- read “What Every Computer Scientist Should Know About Floating Point Arithmetic” conduct a theoretical round-off error analysis
- be aware of de-normalized floating point numbers and their impact on performance
- be aware of non-reproducibility of results when running parallel codes with different core counts
- talk to your local software engineering colleagues
- use static analyzers
- use coverage tools
- use a tool such as `pmccabe` iteratively during code development to keep module complexity low
- use a tool such as Simian to identify duplicated code
- use a tool such as Pylint (http://www.pylint.org/) for improving Python scripts systematically test software
- compare your implementation with other independent implementations

Finally, if we intend to go even one step further, we can consider how software for critical systems such as commercial aircraft autopilots is designed. Typically, a specification is provided to two or three completely independent software development teams. They all develop software that complies with these specifications using different programming languages. Thereafter, given a broad range of input parameters, the outputs of all three independent implementations are compared. This ensures, with high probability, that the autopilot complies with the specification. One must keep in mind though that the specification itself can be incorrect or not cover all cases Thus, in our field, the results of any new tool should be treated with extreme caution until at least one additional, independent implementation is available that yields analogous results. Furthermore, such an independent alternative implementation may also reveal errors in the specification/theory the tool is based upon. An example for this is the detection of an incorrect Hastings ratio calculation for Bayesian inference (Holder et al. 2005) which was unraveled in the course of such an independent implementation effort. We believe that this strategy of comparing the results of independent implementations (e.g., PHYML, IQ-Tree, RAxML for Maximum Likelihood or ExaBayes, MrBayes, PhyloBayes for Bayesian inference) represents a valuable approach to increasing our confidence regarding the correctness of these tools.

In contrast to this, community projects such as R have been very successful, but R also represents a single point of failure. That is, errors in R core modules may have a more dramatic downstream impact than in MrBayes or RAxML, for instance. To this end, we prefer redundancy as *the* mechanism for increasing confidence about correctness.

### Policy Issues

The 15 codes we analyzed have accumulated more than 65,000 citations (not including all papers describing updated versions) based on Google Scholar to date. One may argue that the amount of funding used to generate papers using these codes is disproportional to the amount of funding spent for maintaining and improving these codes, given the catastrophic effects that potential programming or conceptual bugs can have on the published results.

There is a clear lack of sustainable funding for programmers that could maintain and improve the codes developed by PIs or students that leave academia after their PhD. Firstly, one is limited by university or public sector salary schemes which are too low to hire outstanding programmers. Secondly, current funding schemes do not allow for hiring programmers on unlimited time contracts. One option would be to allocate permanent programmer positions to PIs who have an established track record in scientific software development.

One may also consider to allocate temporal funding for re-designing scientific codes to increase maintainability if they rapidly accumulate citations. This could be extended to funding several independent redundant implementations of emerging models and methods. The cost for this is small compared to the potential gains in quality and probability of code correctness.

Another problem is that there is insufficient funding for scientific software development per se. Numerous funding bodies do not consider scientific software development as being ‘real’ research and it is thus extremely hard to obtain financial support. Ironically, a larger number of funded research projects (e.g., a search for the co-occurrence of the terms ‘phylogenetic’ and ‘Deutsche Forschungsgemeinschaft’ yields approximately 17,800 results in Google Scholar) relies on the availability of such tools.

Thus, due to the steadily increasing reliance on computational tools, we believe that novel funding schemes are required to develop new tools as well as improve quality and correctness of existing software. Moreover, the user community must be aware of the fact that, while current tools are freely available, they are developed on a best-effort basis only. There is a plethora of error sources, given that we simply do not have the time nor the resources to implement them properly and occasionally completely re-design them.

Alternatively, one may consider a commercial approach and raise license fees that could be used for providing support and maintenance. One disadvantage of this is that researchers from developing countries may not be able to afford the licenses. In addition, based on our experience with selling non-academic licenses for the PEAR software (Zhang et al. 2014), license management can be time-consuming. Other potential licensing models include crowd-funding, pay-what-you-want strategies, or offering basic, free and advanced, non-free versions of a tool.

## Conclusion

We have presented an initial and simple software quality assessment of widely used evolutionary biology software. We show that by using simple techniques and tools the quality of existing software can already be improved. We also provide a list of best practices for future software development projects. We address issues and provide real-world examples pertaining to numerical reproducibility (or lack thereof) to increase awareness about these issues in the user community. One must also keep in mind that, given the NGS data tsunami, there is a clear trade-off between program performance and maintainability. Programs like RAxML, that explicitly use vector intrinsics for maximum performance on standard laptop/server processor architectures, are substantially harder to maintain. As a consequence of this increased complexity, they are more error-prone than a straightforward naive implementation of Felsenstein’s pruning algorithm.

Further, we argue that the current and rather worrisome state of widely used software in our field is not the fault of the developers, but due to a substantial lack of sustainable funding for software development, improvement, maintenance, and support. This is especially true if one considers the disproportion between funding spent for generating the data with respect to funding spent for improving the quality of software that is being used for analyzing these data. We also make suggestions on how journals, editors, and reviewers could take measures for improving software quality in the course of the review process. Furthermore, the independent development of software by different teams and the comparison of the results can substantially contribute to identifying correctness and not merely quality issues. We are convinced that, in the times of long and complex NGS data analysis pipelines with an ever increasing number of components, software quality issues are becoming critical to the success of the field. Thus, as long as there are no additional efforts on improving software quality, and given the current mediocre quality of our tools, users should not treat evolutionary analysis tools as black boxes, but rather as potential Pandora’s boxes. Apart from improving software quality, we also need to invest more effort into the systematic validation of the results produced by our codes in the future.

## Acknowledgments

We wish to thank Volker Springel, Bastien Bousseau and Tracy Heath for suggestions and discussions regarding this project. We would also like to thank our software engineering colleague Ralf Reussner at KIT for insightful discussions. We are particularly grateful to Mark Holder for extremely useful suggestions and comments on an earlier version of this manuscript. We wish to thank Stephane Guindon and Fredrik Ronquist for their reviews of the initial version of this manuscript. We acknowledge institutional funding by HITS.

## References

Briand, L. C., J. Wüst, J. W. Daly, and D. V. Porter. 2000. Exploring the relationships between design measures and software quality in object-oriented systems. Journal of systems and software 51: 245–273.

Briand, L. C., J. Wüst, S. V. Ikonomovski, and H. Lounis. 1999. Investigating quality factors in object-oriented designs: an industrial case study. Pages 345–354 *in* Proceedings of the 21st international conference on Software engineering ACM.

Drummond, A. J., and A. Rambaut. 2007. BEAST: Bayesian evolutionary analysis by sampling trees. BMC evolutionary biology 7: 214.

Fletcher, W. and Z. Yang. 2009. INDELible: a flexible simulator of biological sequence evolution. Molecular biology and evolution 26: 1879–1888.

Goldberg, D. 1991. What every computer scientist should know about floating point arithmetic. ACM Computing Surveys 23: 5–48.

Grimm, G. W., S. S. Renner, A. Stamatakis, and V. Hemleben. 2006. A nuclear ribosomal DNA phylogeny of acer inferred with maximum likelihood, splits graphs, and motif analysis of 606 sequences. Evolutionary Bioinformatics Online 2: 7.

Guindon, S., J.-F. Dufayard, V. Lefort, M. Anisimova, W. Hordijk, and O. Gascuel. 2010. New algorithms and methods to estimate maximum-likelihood phylogenies: assessing the performance of PhyML 3.0. Systematic biology 59: 307–321.

Heath, T. A., J. P. Huelsenbeck, and T. Stadler. 2014. The fossilized birth-death process for coherent calibration of divergence-time estimates. Proceedings of the National Academy of Sciences 111:E2957-E2966.

Hoare, C. A. R. 1969. An axiomatic basis for computer programming. Communications of the ACM 12: 576–580.

Holder, M. T., P. O. Lewis, D. L. Swofford, and B. Larget. 2005. Hastings ratio of the LOCAL proposal used in Bayesian phylogenetics. Systematic biology 54: 961–965.

Hudson, R. R. 2002. Generating samples under a Wright-Fisher neutral model of genetic variation. Bioinformatics 18: 337–338.

Jarvis, E. D., S. Mirarab, A. J. Aberer, B. Li, P. Houde, C. Li, S. Y. Ho, B. C. Faircloth, B. Nabholz, J. T. Howard, et al. 2014. Whole-genome analyses resolve early branches in the tree of life of modern birds. Science 346: 1320–1331.

Johnson, S. C. 1977. Lint, a C program checker. Citeseer.

Katoh, K. and D. M. Standley. 2013. MAFFT multiple sequence alignment software version 7: improvements in performance and usability. Molecular biology and evolution 30: 772–780.

Li, R., C. Yu, Y. Li, T.-W. Lam, S.-M. Yiu, K. Kristiansen, and J. Wang. 2009. SOAP2: an improved ultrafast tool for short read alignment. Bioinformatics 25: 1966–1967.

Löytynoja, A. and N. Goldman. 2005. An algorithm for progressive multiple alignment of sequences with insertions. Proceedings of the National academy of sciences of the United States of America 102: 10557–10562.

McCabe, T. J. 1976. A complexity measure. Software Engineering, IEEE Transactions on Pages 308–320.

Misof, B., S. Liu, K. Meusemann, R. S. Peters, A. Donath, C. Mayer, P. B. Frandsen, J. Ware, T. Flouri, R. G. Beutel, et al. 2014. Phylogenomics resolves the timing and pattern of insect evolution. Science 346: 763–767.

Myers, G. J., C. Sandler, and T. Badgett. 2011. The art of software testing. John Wiley & Sons.

Nielsen, R., S. Williamson, Y. Kim, M. J. Hubisz, A. G. Clark, and C. Bustamante. 2005. Genomic scans for selective sweeps using SNP data. Genome research 15: 1566–1575.

Notredame, C., D. G. Higgins, and J. Heringa. 2000. T-Coffee: A novel method for fast and accurate multiple sequence alignment. Journal of molecular biology 302: 205–217.

Rambaut, A. and N. C. Grass. 1997. Seq-Gen: an application for the Monte Carlo simulation of DNA sequence evolution along phylogenetic trees. Computer applications in the biosciences: CABIOS 13: 235–238.

Robinson, D. and L. Foulds. 1981. Comparison of phylogenetic trees. Mathematical Biosciences 53:131–147.

Ronquist, F., M. Teslenko, P. van der Mark, D. L. Ayres, A. Darling, S. Hhna, B. Larget, L. Liu, M. A. Suchard, and J. P. Huelsenbeck. 2012. MrBayes 3.2: Efficient Bayesian phylogenetic inference and model choice across a large model space. Systematic Biology 61: 539–542.

Simpson, J. T., K. Wong, S. D. Jackman, J. E. Schein, S. J. Jones, and I. Birol. 2009. ABySS: a parallel assembler for short read sequence data. Genome research 19: 1117–1123.

Springel, V. 2005. The cosmological simulation code gadget-2. Monthly Notices of the Royal Astronomical Society 364: 1105–1134.

Yang, Z. 2007. PAML 4: phylogenetic analysis by maximum likelihood. Molecular biology and evolution 24: 1586–1591.

Yang, Z. and B. Rannala. 2010. Bayesian species delimitation using multilocus sequence data. Proceedings of the National Academy of Sciences 107: 9264–9269.

Zhang, J., K. Kobert, T. Flouri, and A. Stamatakis. 2014. Pear: a fast and accurate illumina paired-end read merger. Bioinformatics 30: 614–620.

